# The SCWISh network is essential for survival under mechanical pressure

**DOI:** 10.1101/150789

**Authors:** Morgan Delarue, Gregory Poterewicz, Ori Hoxha, Jessica Choi, Wonjung Yoo, Jona Kayser, Liam Holt, Oskar Hallatschek

## Abstract

Cells that proliferate within a confined environment build up mechanical compressive stress. For example, mechanical pressure emerges in the naturally space-limited tumor environment. However, little is known about how cells sense and respond to mechanical compression. We developed microfluidic bioreactors to enable the investigation of the effects of compressive stress on the growth of the genetically tractable model organism Saccharomyces cerevisiae. We used this system to determine that compressive stress is partly partly sensed through a module consisting of the mucin Msb2, and the cell wall protein Sho1, which act together as a sensor module in one of the two major osmosensing pathways in budding yeast. This signal is transmitted via the MAPKKK kinase Ste11. Thus, we term this mechanosensitive pathway the *SMuSh* pathway, for *S*te11 through *Mu*cin / *Sh*o1 pathway. The SMuSh pathway delays cells in the G1 phase of the cell cycle and improves cell survival in response to growth-induced pressure. We also found that the Cell Wall Integrity (CWI) pathway contributes to the response to mechanical compressive stress. These latter results are confirmed in complimentary experiments in the accompanying manuscript from Mishra et al. When both the SMuSh and the CWI pathways are deleted, cells fail to adapt to compressive stress and all cells lyse at relatively low pressure when grown in confinement. Thus, we define a network that is essential for cell survival during growth under pressure. We term this new mechanosensory system the SCWISh (Survival through the CWI and SMuSh) network.

**Significance Statement:** Growth in confined environments leads to the build up of compressive mechanical stresses, which are relevant to diverse fields, from cancer to microbiology. In contrast to tensile stress, little is known about the molecular integration of compressive stresses. In this study, we elucidate the SMuSh pathway, which, together with the Cell Wall Integrity pathway, is essential for viability of the budding yeast *S. cerevisiae* when growing under mechanical pressure. Pressure-sensing requires the transmembrane mucin, Msb2, which is linked to the actin cortex. Our result raises the intriguing question of whether mucins, widely conserved in eukaryotes and frequently misregulated in cancers, might sense compressive stresses in other organisms, including humans.

---

Mechanical stresses can broadly be separated into stresses of opposite signs: tensile stresses that arise from stretching cells, and compressive stresses that arise from pushing on cells. Tension naturally emerges from cell-cell and cell-substrate adhesion, and has been shown to affect cell proliferation, cell death, cell migration, and even cell differentiation(1–5). In contrast, compressive mechanical stresses often arise due to proliferation in a spatially confined environment, and do not require cell adhesion. Pressure can develop in many types of cell, from microbes(6) to mammalian cells, both in the context of normal tissues and in cancer(7–9).

Tensile stresses have been widely investigated, and much is known about the molecular integration of these stresses (10). In contrast, relatively little is known about the molecules that detect and react to growth-induced compressive mechanical stress. While tensile stresses are restricted to cells that adhere to their environment through molecules connected to their contractile cytoskeleton, such as mammalian cells, compressive stresses can be experienced by any cell population. Recent experiments suggest that the fungus *S. cerevisiae* senses and adapts to compressive mechanical stress(6). To reveal the molecular basis of this mechanosensing, we developed new microfluidic devices to study the effect of growth-induced compressive stress on *S. cerevisiae*, which we discovered employs elements of osmosensing pathways, namely the Msb2 / Sho1 module. This module signals to Ste11, which in turn activates the HOG (high osmolarity glycerol) pathway. We unraveled a mechanosensing pathway that we term *SMuSh*: pressure activates the MAPKKK *S*te11 through the *Mu*cin Msb2 / *Sh*o1. We also found, as in the accompanying manuscript from Mishra et al, that *S. cerevisiae* also employs elements of the Cell Wall Integrity (CWI) pathway to adapt and survive to compressive stress. Deletion of both SMuSh and CWI pathways lead to a total loss of viability when cells develop compressive mechanical stress through growth in confinement. Therefore, we define the SCWISh network (Survival through the CWI and SMuSh), as a new mechanosensory system that is essential for the survival of yeast cells growing under pressure.

## Results

We developed a microfluidic platform to study the growth of *S. cerevisiae* cells under a defined compressive stress. We improved the design of a previously developed confining microfluidic device to enable the imaging of a larger number of cells in an easier-to-handle device(6) (SI Appendix, Fig. S1). Unconstrained cell proliferation occurred in the chamber until cells filled it, at which point further proliferation resulted in the progressive build-up of growth-induced pressure with a typical timescale of ~10h (Fig. 1A and Movie S1). A set of narrow channels enabled the flow of culture medium from the side of the chamber thus supplying nutrients and maintaining a constant chemical environment. Pressure was calculated by quantifying the deformation of the PDMS walls of the chamber (SI Appendix, Fig. S2). We chose a pressure of ~0.4 MPa as the set-point for our analysis, which is about half of the pressure at which wild-type cells stall growth(6). Importantly, all mutant strains were able to generate this pressure, thus enabling direct phenotypic comparison.

**Fig. 1.**
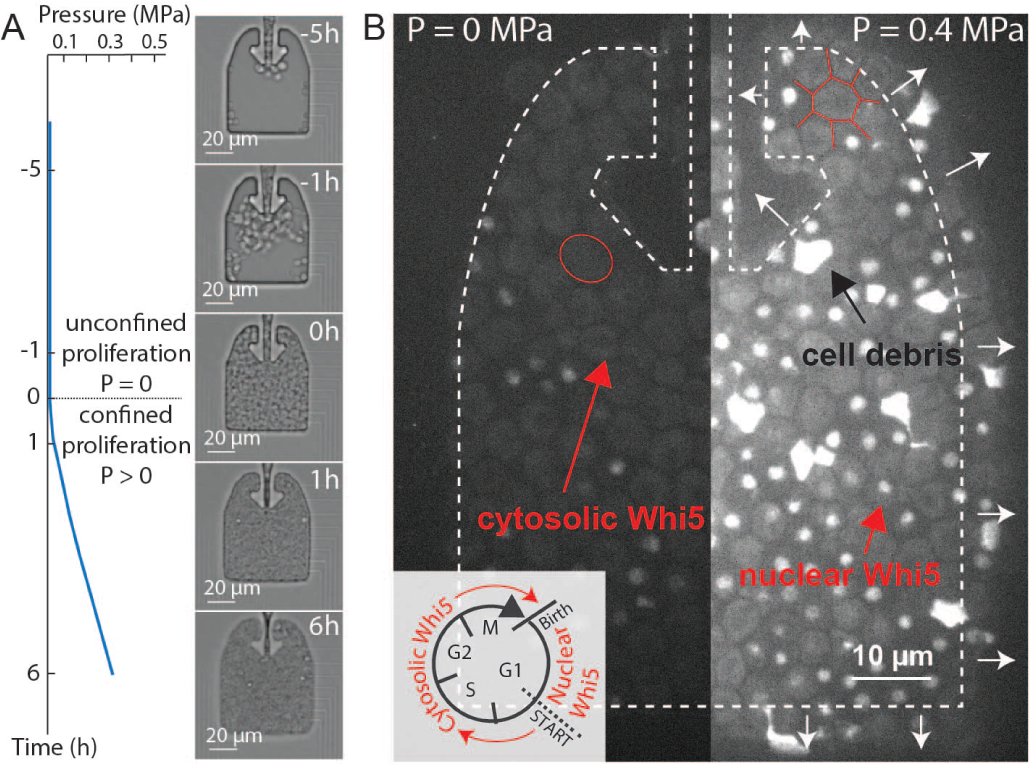
Growth-induced mechanical pressure leads to cell cycle arrest in G1. A. Cells initially proliferated unconfined, developing no pressure, until they fill the confining chamber. At this point, proliferation led to build-up of compressive pressure within hours. We calculated the pressure developed by the cells through the deformation of the PDMS chamber (SI Appendix, Fig. S2). **B**. Nuclear accumulation of *WHI5-mCherry* indicates a delay in the G1 phase of the cell cycle. The contour of the chamber is outlined in dash, white arrows indicating chamber deformation. Red arrows point to cytosolic Whi5 and nuclear Whi5, while the black arrow points to a cell debris. The outline of a typical non-deformed (P = 0 MPa) and a typical deformed (P = 0.4 MPa) cell is shown in red.

In contrast to osmotic stress, which causes isotropic reduction of cell volume without major shape changes(11), cell-cell contact forces imposed by growth in confinement led to severe cell deformation (Fig. 1B). As compressive stress built up, the average cell size was reduced (SI Appendix, Fig. S3), and the rate of proliferation was decreased. Using a *WHI5-mCherry* strain, we found that cells were progressively more delayed in G1 as pressure built up (Fig. 1B) (6), as indicated by a nuclear Whi5 signal(12). We also noticed that about 10% of the cell population died when grown to 0.4 MPa of pressure, as evidenced by accumulation of autofluorescent cell debris (Fig. 1B, Fig 2A). Thus, even our wild-type control cells suffer a significant degree of mortality at 0.4 MPa of pressure, suggesting that this type of stress presents a significant risk to cells.

**Fig. 2.**
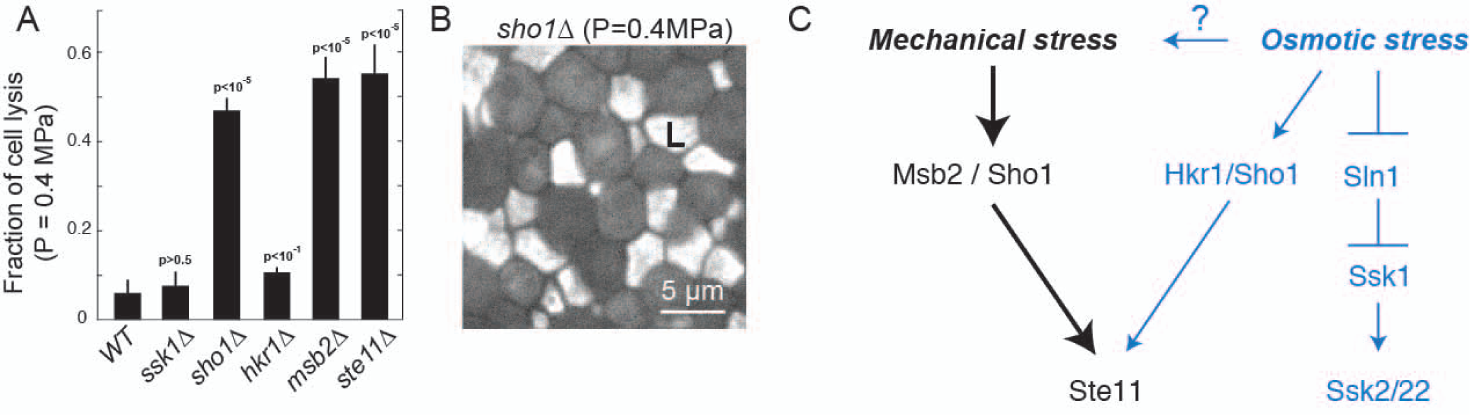
The SMuSh pathway is necessary for survival under compressive mechanical stress. **A**. Fraction of cell lysis at a mechanical pressure of 0.4 MPa in different genetic backgrounds. p-values were calculated by a T-test on more than 5 replicates for each genetic background compared to wild-type (*WT*, more than 100 cells per replicate). **B.** Fluorescent picture of a *sho1*Δ background under mechanical stress, displaying the accumulation of cell debris after lysis. The annotated “L” stand for a cell that underwentlysis under pressure. **C**. Pathway diagrams for putative osmosensors in *S. cerevisiae*. Our results suggest that the Msb2/Sho1 module is a mechanosensor (black) while Sln1 and Hkr1/Sho1 modules are osmosensors (blue).

The progressive enrichment of cells in the G1 phase of the cell cycle, together with the occurrence of cell death suggested that inhibition of proliferation under mechanical stress could be an adaptation to increase survival in this challenging environment. This model implies the existence of molecular pathways that sense and respond to compressive stress. We hypothesized that this mechanosensing could employ elements of the osmosensing machinery because both stresses result in cell volume reduction and water efflux.

In budding yeast, two overlapping osmosensing branches have been identified, both of which activate MAP kinase (MAPK) cascades(11). The SLN1 branch regulates the activity of the MAPKKKs Ssk2 and Ssk22 under osmotic stress, whereas the SHO1 branch activates the MAPKKK Ste11. Both branches converge to activate the MAPK Hog1, which is thought to be the primary effector of the osmotic stress response.

We explored whether genetic alterations to these branches would lead to differential cell survival in response to mechanical and osmotic stress. We found that, while both branches respond to and promote cell survival under osmotic stress (SI Appendix, Fig. S4), disrupting the *SLN1* branch by deletion of *SSK1* did not affect cell survival under mechanical stress (Fig. 2A), suggesting that the *SLN1* branch is dispensable for survival under compressive mechanical stress. In clear contrast, deletion of *SHO1* led to dramatic cell death: close to 50% of cells died when the population reached a pressure of 0.4 MPa (Fig. 2A-B). Thus, the *SHO1* branch is required for survival during growth-induced compressive stress. Note that the presence of cell debris, albeit mechanically different from living cells, do not chemically influence cell proliferation (see Methods)

The Sho1 protein has two different mucin co-activators: Msb2p and Hkr1p. These are high molecular weight, membrane bound glycoproteins that communicate with Sho1p through a poorly understood mechanism(13, 14). Deletion of the *HKR1* mucin only had a mild effect on cell survival, but deletion of *MSB2* caused a dramatic cell death phenotype under mechanical stress as *SHO1* deletion (Fig. 2A). This genetic result is distinct from that reported using zymolyase treatment. *HKR1* is required for survival in the zymolyase model cell wall stress, while MSB2 is dispensable in this context(15). Deletion of the *STE11* kinase, which is downstream of Sho1, also led to dramatic cell death under pressure. Together, these results suggest that MSB2/SHO1 senses compressive mechanical stress and activates the MAPKKK Ste11 (Fig. 2C). We chose to term this pathway the SMuSh pathway, for the activation of *S*te11 through *Mu*cin Msb2/*Sh*o1.

We sought to determine if cells that lack components of the SMuSh pathway are intrinsically unstable when mechanically compressed. To address this question, we developed a new microfluidic device that allowed us to instantaneously exert mechanical compressive stress (Fig. 3A). In this system, cells were loaded into a growth chamber and then confined by sealing the input/output valve. Subsequently, pressure was induced in one of two alternative ways: Either the confined cells were allowed to divide to build-up growth-induced pressure over several hours (Fig. 3B and SI Appendix, Fig. S5), or a thin membrane “micro-piston” at the base of the chamber was distorted to instantaneously compress the cell population (Fig. 3C). When cells progressively built up pressure through growth and division there was a large increase in cell death in the *ste11*Δ background compared to wild-type cells (Fig. 3D). However, when cells were instantaneously compressed to a comparable pressure, there was no cell death in either strain (Fig. 3E). These results demonstrate that loss of SMuSh components does not cause intrinsic mechanical instability. Rather, the cell death phenotype that we observe in mutants for the mechanosensing pathway only occurs when cells grow and/or divide under pressure.

**Fig. 3.**
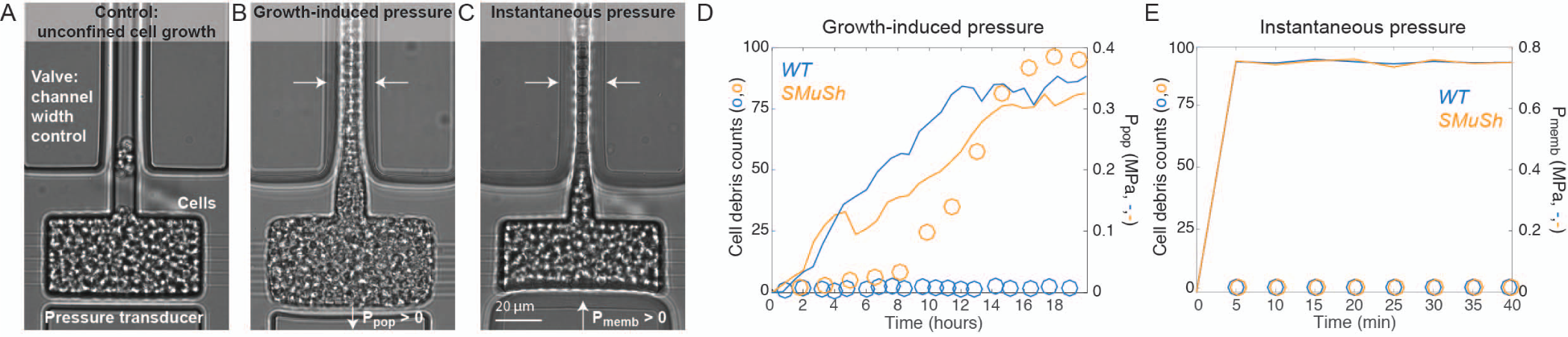
SMuSh pathway mutants mainly lyse during S/M phase and are not intrinsically mechanically unstable. When the input/output valve is open (**A**), cells can readily flow out of the chamber and therefore do not build-up a mechanical stress. When the valve is closed, cells are confined and eventually develop growth-induced mechanical pressure (**B**). Alternatively, cells can be instantaneously compressed using a pressure transducing micropiston (**C**). We observed an accumulation of cell debris under the build-up of pressure in a *ste11*Δ background but not in the wild-type (**D**). Instantaneous compression does not increased cell death in either strain, suggesting that the cells are not intrinsically unstable, but only lyse if they attempt to grow under pressure (**E**).

We used both the subcellular localization of Whi5 in *WHI5mCherry* cells and the spindle shape in *GFP-TUB1* cells to assess the position of cells in the cell cycle. The subcellular localization of Whi5 depends on CDK activity and the shape of spindle gives an additional indication of cell cycle phase (16). Examples of cells in which the spindles in different phases of the cell cycle are displayed in SI Appendix, Fig. S6. In wild-type cells, the build-up of compressive, growth-induced mechanical stress was accompanied by an increase in cells delayed in the G1 phase of the cell cycle, (Fig. 4A-B). We observed that this cell cycle delay was abrogated in strains deleted for SMuSh components (Fig. 4A-B, note the large increase of cells in S/M phases in SMuSh mutants). Therefore, we hypothesized that cell cycle arrest was important for cell survival. In agreement with this idea, all *ste11*Δ cells that we saw lysing (N ≥ 10) had a cytosolic Whi5 signal prior to death, indicating that these cells had progressed beyond START to enter the cell division cycle (SI Appendix, Fig. S7). Using GFP-TUB1, we found that the great majority of cell lysis occurred during S/M phase (Fig. 4D). We speculate that the polarized growth that drives budding in S and M phase creates a mechanical instability that makes *S. cerevisiae* prone to lysis under pressure.

**Fig. 4.**
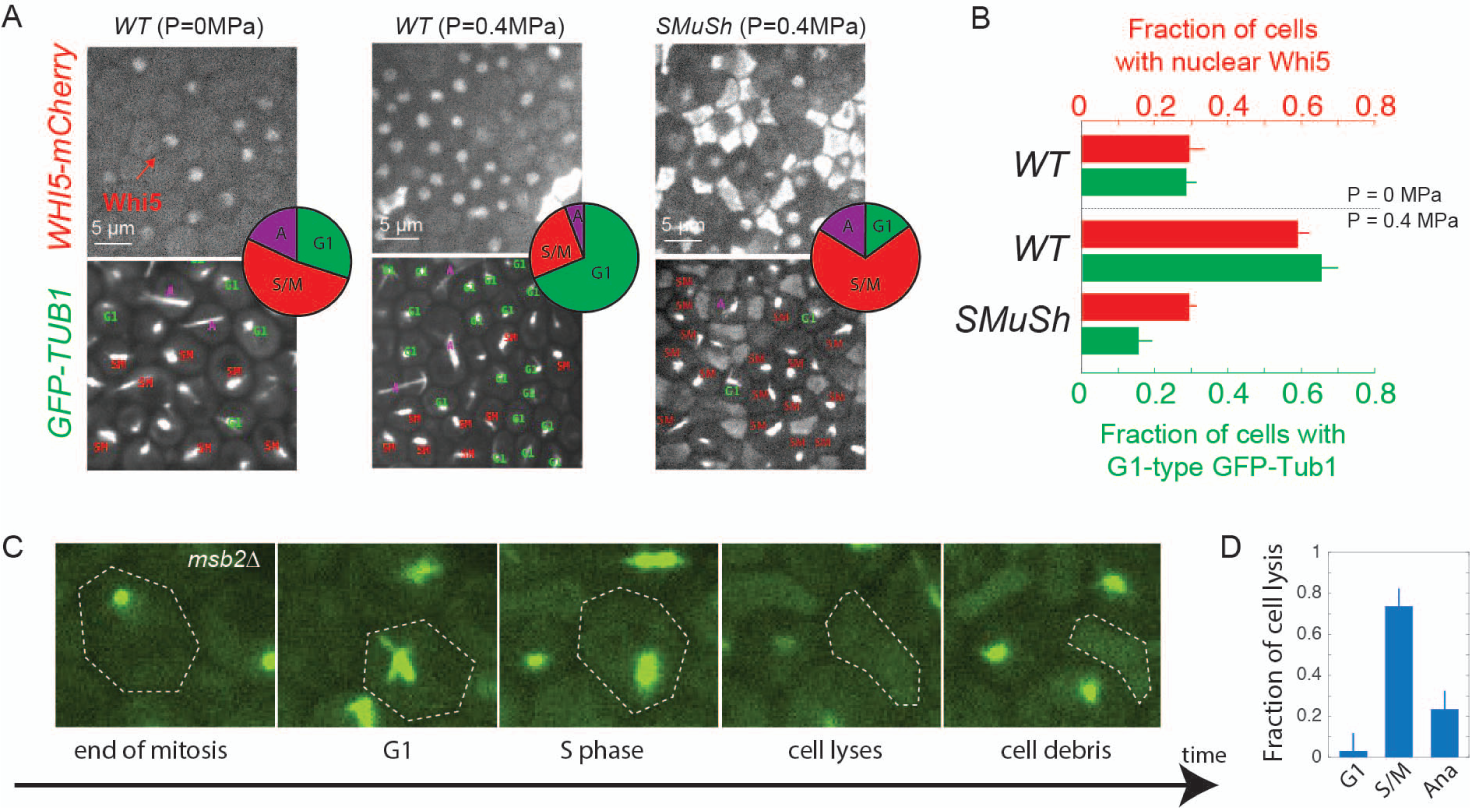
The SMuSh pathway delays cells in the G1 phase of the cell cycle. **A**. Increased compressive stress results in the nuclear retention of Whi5-mCherry and accumulation of cells displaying with G1 asters in *GFP-TUB1* cells. Deletion of the SMuSh pathways results in extensive cell lysis and a reduction in the fraction of cells in G1. Inset: Pie charts showing the fraction of cells in various phases of the cell cycle as determined from GFP-Tub1 and cell morphology. **B**. Quantification of the fraction of cells in G1, through *WHI5-mCherry* localization, GFP-Tub1 and cell morphology. Analysis of movies of GFP-TUB1 strains, revealed that cells mainly lyse during the S and M phases of the cell cycle (**C**).

Activation of Ste11 has been reported to activate two main pathways: the osmotic response pathway, through the Mitogen Activated Protein Kinase (MAPK) Hog1 and the invasive growth pathway, through the MAPK Kss1(17). In addition, Ste11 has recently been shown to signal to the cell wall integrity pathway and its MAPK Slt2 (15, 18–21)(Fig. 5A). When Hog1 is activated, it undergoes a translocation from the cytoplasm to the nucleus. We observed nuclear relocalization of the MAPK Hog1 both under instantaneous and growth-induced compressive stress suggesting this MAPK is responsive to both stresses (Fig. 5B). The degree of nuclear translocation in response to a 0.4 MPa mechanical compressive stress was comparable to that observed with a 1M osmotic stress (Fig. 5C). As expected from our results above, deletion of *SHO1* resulted in a significant decrease in the fraction of cells that activate Hog1. In contrast, disruption of the *SLN1* branch by deletion of *SSK1* had not effect. These results indicate that compressive and osmotic stresses both activate Hog1, but that mechanical pressure predominantly uses the Sho1/Msb2 sensor.

**Fig. 5.**
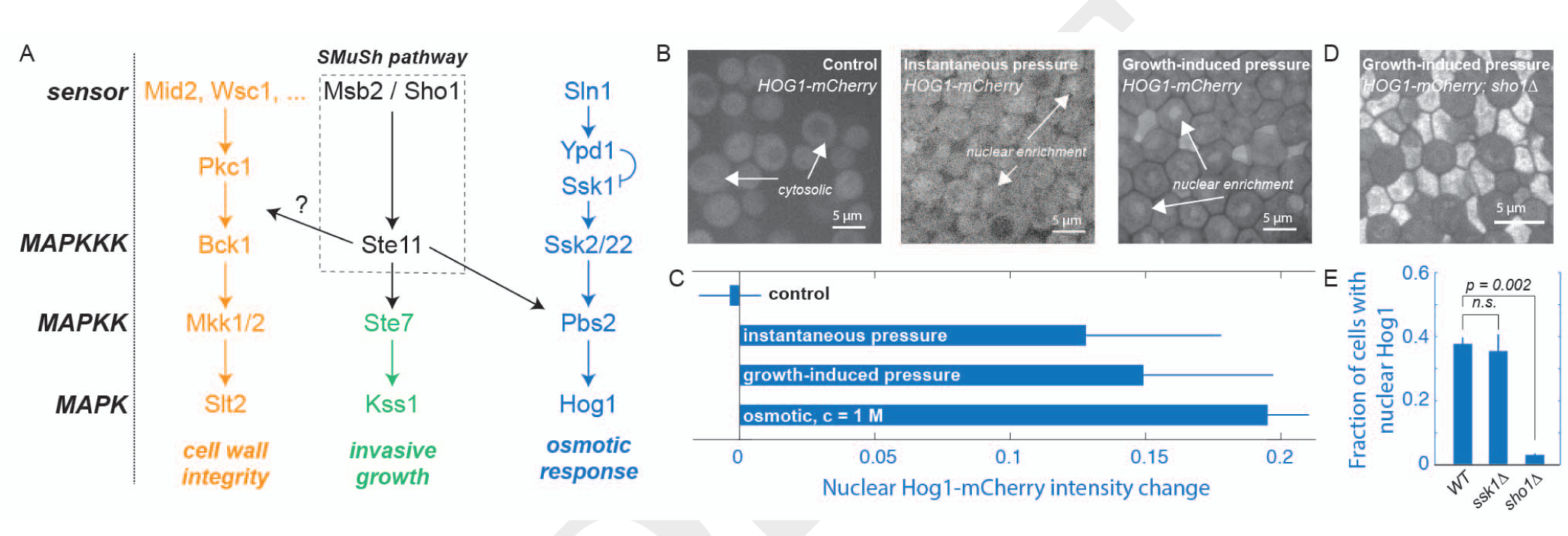
The HOG pathway is activated in response to compressive stress in a SMuSh-dependent manner. **A**. The upstream kinase Ste11 drives the invasive growth and HOG MAP kinase pathways through the downstream MAP kinases (MAPKK/MAPK) Ste7/Kss1, and Pbs2/Hog1 respectively. Both instantaneous and a growth-induced compressive stress result in the HOG pathway activation as determined by nuclear translocation of Hog1-GFP (**C**). Deletion of SMuSh components such as *sho1*Δ led to a large increase in cell lysis (**D**) and a significant decrease of fraction of cells that activated the HOG pathway (**E**).

Single deletion of the MAPKs Kss1 or Hog1 only mildly increased the fraction of cell lysis (Fig. 6D). In contrast, *kss1*Δ;*hog1*Δ double mutant strains suffered significant (60%) cell lysis at 0.4 MPa pressure, a phenotype comparable to that of SMuSh pathway deletion strains. This result strongly suggests that the Kss1 and Hog1 MAPKs act synergistically to enable cells to adapt to mechanical compression.

**Fig. 6.**
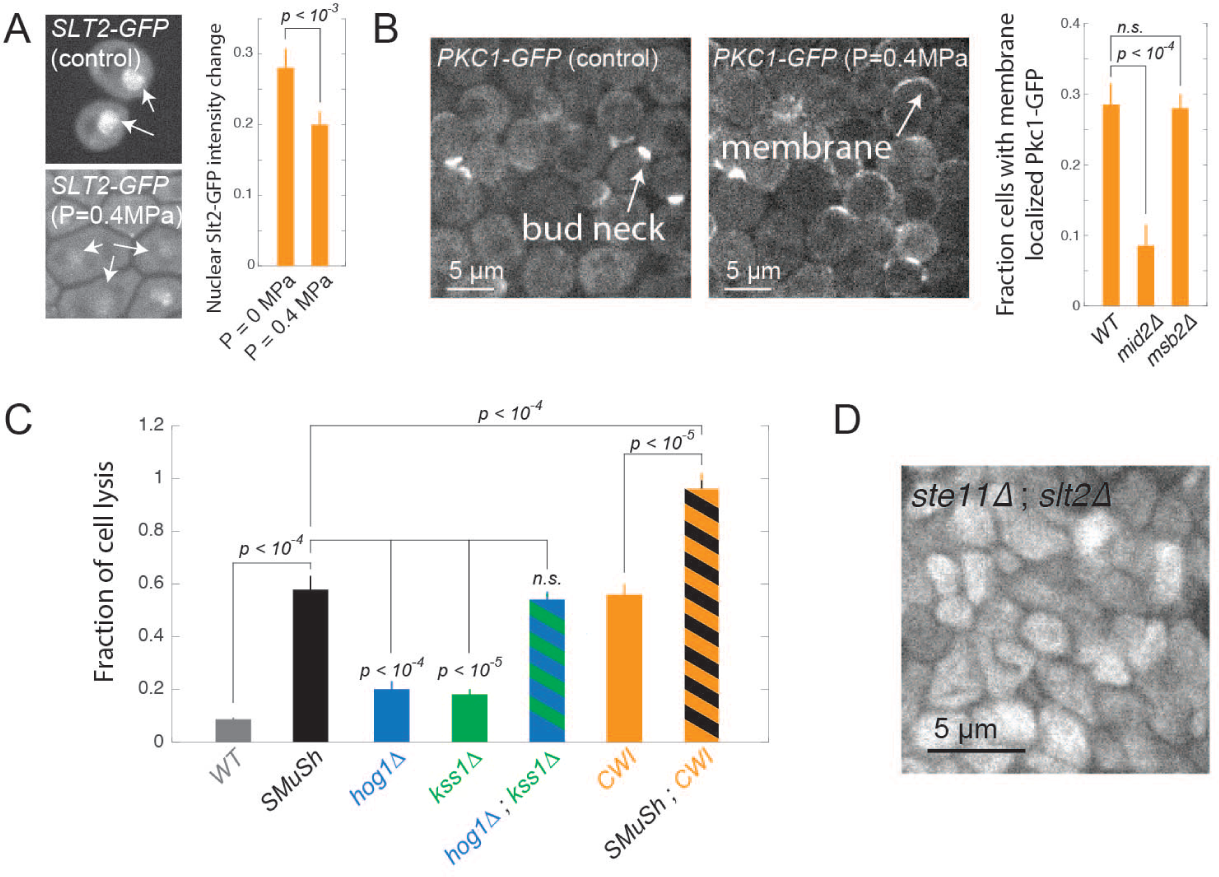
The SCWISh network, which includes both SMuSh and the CWI pathways, is essential for survival during growth under pressure. **A**. Slt2-GFP translocates from the nucleus to the cytoplasm in response to compressive mechanical stress. **B**. Pkc1p is recruited from the cytoplasm to the plasma membrane in response to compressive mechanical stress. Deletion of the mucin Msb2p of the SMuSh pathway had no effect on CWI pathway activation. **C**. Deletion the Kss1 or Hog1 MAPKs individually led to a mild cell lysis phenotype, but the double delete *kss1*Δ;*hog1*Δ had a comparable degree of cell lysis as mutants in the upstream SMuSh pathway components (*MSB2*, *SHO1* and *STE11*). Simultaneous abrogation of the SMuSh and CWI pathways, in double mutant *ste11*Δ;*slt2*Δ cells led to a complete loss of viability under compressive mechanical stress. (**D**).

We then asked whether SMuSh could activate the CWI pathway. The CWI pathway has been reported to be important for viability in a wide variety of stress conditions (15, 18–21) and so we reasoned that CWI may play a role the response to mechanical pressure. In contrast to Hog1, the effector MAPK of the CWI pathway, Slt2, translocates from the nucleus to the cytoplasm when activated. In addition, Protein kinase C (Pkc1), another key kinase in this pathway, is recruited to the plasma membrane when activated. We observed both nuclear delocalization of Slt2 and plasma membrane relocalization of Pkc1 in response to compressive stress (Fig. 6A). Together, these results indicated that the CWI pathway was indeed activated by mechanical pressure (Fig. 6B). The importance of the CWI pathway during compressive stress is also reported in the accompanying paper in this issue of PNAS (22). Deletion of the CWI membrane sensor, *MID2*, significantly decreased the fraction of cells that relocalized Pkc1-GFP (Fig. 6B) and led to a 30% increase in cell death (SI Appendix, Fig. S8). A double *mid2*Δ;*slt2*Δ yielded the same fraction of cell lysis as a single *slt2*Δ, suggesting that these genes are epistatic and that Mid2 and Slt2 are in the same pathway, but more than *mid2*Δ alone, suggesting that compressive stress may activate Slt2p through other types of sensors (SI Appendix, Fig. S8). In contrast, deletion of the SMuSh component *MSB2* did not affect Pkc1 relocalization, suggesting that the SMuSh pathway does not act through the CWI pathway but rather is a parallel pathway in a network that responds to compressive stress. Deletion of the MAPK Slt2p significantly increased the fraction of cell lysis under pressure (Fig. 6C), as did a deletion of the CWI MAPKKK Bck1p (SI Appendix, Fig. S8), further suggesting that activation of CWI is necessary for cells to adapt and survive during growth under compressive mechanical stress.

To test for synthetic lethality and determine if SMuSh and CWI impinge upon the same process, we deleted both *STE11* and *SLT2*. These double mutants underwent a dramatic and complete cell lysis even at relatively low pressure (≤ 0.2 MPa) (Fig. 6C-D). This result shows that the SMuSh and CWI act together to allow adaptation to pressure. Thus, we have defined a new mechanosensitive system which we term the **SCWISh** (**S**urvival through **CWI** and **S**MuS**h**) network, essential for cell survival during growth under compressive mechanical stress.

## Discussion

Compressive mechanical stress can affect any cell population that proliferates in a space-limited environment. However, until now, the study of compressive mechanical stress has been limited to a few examples in mammalian cells (7–9) owing to the experimental difficulty of imposing a controlled compressive stress while keeping the chemical environment constant. In this paper, we present two new microfluidic devices that enabled us to study the effects of growth induced on the genetically tractable model *S. cerevisiae*. Using these devices, we were able to apply either growth-induced or instantaneous compressive stress (6). The former enabled the study of adaptation to compressive stress, whereas the latter enabled us to study the acute effects of compressive stress.

Using these devices, we found that survival under compressive mechanical stress requires two pathways: The SMuSh pathway, which appears to employ the Kss1 and Hog1 MAP kinases as its effectors, and the CWI pathway, through a set of effectors including the Pkc1, Bck1 and Slt2 kinases. Thus, our work, together with the accompanying paper from Mishra et al (22), defines the first system essential for survival during growth under pressure pressure. We term this system the SCWISh network.

The molecular details of how compressive mechanical stress is sensed remain to be determined in future work. However, we can speculate on various sensing possibilities. The SMuSh pathway is connected to the actin cytoskeleton via the transmembrane mucin Msb2p. We have found that compressive stress can strongly deform the cells. This deformation could affect the cytoskeleton or lead to changes in membrane tension leading to the activation of the pathway by similar mechanisms to those known to operate in response to tensile stress (10). Alternatively, cells could sense the compression of the periplasm (the space between the plasma membrane and the cell wall). It has been found that the highly glycosylated part of Msb2p is necessary for its activity (23), and the CWI upstream sensors such as Mid2p may be also sense the dimensions of the periplasm (24).

SCWISh mutant cells grow slowly but divide with relatively low rates of cell lysis in the absence of compressive stress. However, spontaneous cell death is observed during < 5% of divisions (SI Appendix, Fig. S9). Note that we did not observe a significant increase in cell death for this mutant when exposed to an osmotic stress, suggesting that the synthetic lethality of a SCWISh mutant is specific to mechanical stress (SI Appendix, Fig. S9).

Further studies will also be required to elucidate the adaptive response downstream of the SCWISh network. However, we do have several clues from the literature. For instance, progression beyond START is controlled by Hog1p through stabilization of the Cyclin Dependent Kinase 1 inhibitor, Sic1p (25) and the initiation of DNA replication is inhibited by Slt2p through degradation of the pre-replicative complex component Cdc6 (26). We speculate that the SCWISh network leads to delays or arrests at multiple cell cycle checkpoints to prevent cell growth and division when cells are exposed to the mechanical challenges of compressive stress.

Our results identified the *SMuSh* pathway required for cell survival in mechanical compressive stress. The identification of the Msb2 / Sho1 / Ste11 module as the key sensor in this mechanosensing paradigm opens new avenues to understand the physical details of compressive mechanosensing. This core SMuSh pathway seems to be conserved in various pathogenic fungi (27–30). Transmembrane mucins are also important in human physiology and are frequently overexpressed in cancer(31, 32), another context where compressive stresses arise from local cell proliferation (8, 9). In this context, the level of mechanical stress is more in the kPa range, because mammalian cells do not possess a cell wall. However, we have seen that a 5 KPa compressive stress can lead to biophysical changes in mammalian cells (33) that require 0.4 MPa of stress in *S. cerevisiae*. Thus, we suggest that similar pathways could operate in mammalian cells but be adapted to a very different pressure range. Our results raise the intriguing possibility that mechanosensing through mucins may be widely conserved in eukaryotes, with or without a cell wall.

## Materials and Methods

### Yeast culture and transformation

All cells were cultured in synthetic complete (SC) medium supplemented with 20g/L of dextrose (D), at 30°C. Exponentially growing cells in SCD were loaded at O.D. ~0.3 in the microfluidic device, and fed with SCD at 30°C. All strains used in this study are isogenic to the BY4741 background (SI Appendix, Table S1), obtained from the Yeast Deletion Collection. Yeast strains were created by transforming with a LiAc based approach according to standard methods.

### Imaging

Every condition was imaged with a Nikon TI-Eclipse spinning-disk microscope at 100x magnification, and the images recorded with a scMOS camera (Zyla, Andor). mCherry-tagged proteins were imaged using 560 nm illumination and strains expressing GFP-tagged proteins were imaged at 480 nm.

### Microfabrication and microfluidic device preparation

The mold consists of 2 layers of different heights, each layer prepared using a classical soft lithography protocol described in Ref. (34). The first layer is prepared using SU 2000.5 negative photoresist (0.5 *¼*m height), and the second using SU 2010 (10 *¼*m height). Polydimethylsiloxane (PDMS, Sylgard 184, Dow Corning, USA) is mixed with the curing agent (ratio 1:10 in mass), poured onto the mold, and cured overnight at 60°C. PDMS is bound to glass slides through an oxygen plasma generated by a reactive ion etcher (RIE) machine (P_02_ = 200 mTorr, exposure time = 20 sec).

### Quantification of the fraction of cell debris and nuclear Whi5

We left cells to develop a growth-induced pressure to a typical value of 0.40 ± 0.05, and compared physiological changes to a situation at the beginning of the experiment, or just before the chamber was filled and the cells were about to develop pressure. For each condition, a fixed 50*¼*m × 50*¼*m region of interest, containing about 100 cells, was extracted from the fluorescent image. Within this region, we counted the total number of cells, the number of cell debris, and, when relevant, the number of cells displaying a Whi5 nuclear signal. The fraction of cell debris was calculated relative to total cell number, and the fraction of cells displaying a Whi5 nuclear signal was calculated relative to the number of live non-cell debris for at least 5 biological replicates for each condition. A Student’s T-test (ttest2 function in Matlab) was used to calculate the p-value. A difference was considered non-significant (n.s.) when the p-value was larger than 0.1, otherwise, the p-values are displayed on the figures. We artificially created cell debris by crushing cells in a cryogenic mill ball, and mixed a mass of 50% of cell debris to wild type cells, in order to assess the chemical effect of the presence of cell debris on cell growth. This led to a relative growth rate difference of 2 ± 4%, suggesting no effect of the presence of cell debris on growth rate.

## ACKNOWLEDGMENTS

The authors would like to thank J. Hartung for his initial help with the microfabrication and experiments/ simulations, as well as the Thorner and Boeke labs for some of the strains used in this study. Research reported in this publication was supported by the National Institute Of General Medical Sciences of the National Institutes of Health under Award Number R01GM115851 (O.H.). The content is solely the responsibility of the authors and does not necessarily represent the official views of the National Institutes of Health.

